# Universal donor plasmids for biallelic degron-tagging in mammalian cells and organoids

**DOI:** 10.64898/2026.09.14.751494

**Authors:** Sebastian Öther-Gee Pohl, Brianda A Hernández Morán, Camila Guzman Cardozo, Gillian CA Taylor, Abram Giller, Kevin B Myant, Sara Macias, Andrew J Wood

**Affiliations:** Institute of Genetics and Cancer, University of Edinburgh, Edinburgh, UK; Institute of Immunology and Infection Research, University of Edinburgh, Edinburgh, UK

## Abstract

Degron tags enable rapid, tuneable and reversible control of protein expression, and have become a critical tool for target validation and elucidating biological mechanisms. Tag engineering uses genome editing to generate biallelic gene fusions, though this process remains laborious and requires new donor vectors to be generated for each locus. Here, we have generated a series of universal donor vectors to enable biallelic integration of different degron tags (AID, dTAG, BromoTag, SMASh) using different selection strategies (FACS, drug selection). Using this system, tags can be fused in-frame to endogenous protein C-termini via homology-independent repair following Cas9-induced double strand breaks, as shown here using human lymphoma cells, mouse embryonic stem cells and colonic tumour organoids. This workflow simplifies the genetic engineering of degron tags and is applicable in contexts where homology-dependent repair is challenging.

## Introduction

Targeted protein degradation enables specific proteins to be rapidly destroyed in response to small molecule ligands ^1^. These ligands, known collectively as ‘degraders’, often work by inducing proximity between the target protein and an E3 ubiquitin ligase, leading to ubiquitination of the target followed by proteasomal degradation. Over the past decade, this principle has simultaneously provided new approaches for drug development and new methods to study fundamental cellular processes. In experimental studies, gene knockout or knockdown approaches often need many hours or days to take effect, whereas protein degradation can proceed over tens of minutes. This provides temporal resolution to understand the consequences of perturbation at acute timescales, which is particularly important when studying fast processes like signalling or the cell cycle, or when chronic perturbation induces lethality ^2–4^. Degrader titration and washout also provide straightforward methods for perturbations to be tuned and reversed.

Although an increasing number of degraders have been developed to directly target endogenous proteins ^5^, a large majority of potential targets lack suitable ligands. In these cases, genetic engineering can be used to fuse a “degron tag”, encoding an inducible E3 ligase binding domain with activity that can be regulated using a generic degrader ligand ^6,7^. Several classes of inducible degron tag have been developed. These include the dTAG and BromoTag systems, which use bifunctional PROTAC inducers that simultaneously engage target and E3 ligase using distinct chemical groups^4,8,9^. Other systems such as the auxin-inducible degron (AID/AID2) use monovalent ‘molecular glue’ ligands that first bind to an E3 ligase subunit and change the surface structure to facilitate degron peptide binding ^10–12^. The Small Molecule Assisted Shutoff (SMASh) system works via a third mechanism where the tag encodes a self-cleaving hydrophobic degron domain ^13^. Induction is achieved via chemical inhibition of the cleavage reaction to generate an unstable tagged protein.

Regardless of the system being used, fusion protein expression requires genomic insertion of sequences encoding the degron tag in-frame with the endogenous gene. Insertions are targeted by combining Cas9-mediated double strand breaks (DSBs) with donor vectors containing stretches of homology to flanking DNA sequences ^14,15^. Consequently, fusions at different genes, or of different tags at the same gene, require new donor vectors to be constructed. The biochemical pathways underlying homology-dependent DNA repair are also highly variable across cell types and cell cycle phases ^16^, so genetic approaches relying on homology-dependent repair pathways are not always viable.

Alternative approaches for tag engineering achieve genomic integration without homologous donor sequences by harnessing the non-homologous end joining pathway to repair DSBs targeted by Cas9 ^17–20^. In particular, the CRISPaint system was developed for tagging with fluorescent proteins and uses two sgRNAs: a target selector and a frame selector. CRISPaint donor vectors contain three “frame selector” spacer sequences (0, +1, or +2) which are offset by a single nucleotide. A target selector sgRNA is chosen first to cut upstream from the stop codon of the target gene, then the appropriate frame selector is chosen to linearise the donor in a way that matches the reading frame of the target gene at the Cas9 cut site ^19^.

Despite this clever approach to enrich desirable edits, gene tagging remains inefficient and cells carrying the desired edits are typically rare. In the case of degron tagging, this issue is compounded by the requirement to obtain cells carrying tag fusions on both alleles to ensure the full protein pool is sensitive to degrader induction. For HDR-based approaches, this issue has been addressed by combining orthogonal markers for selection of biallelic insertions ^14,21^. Here, we have adapted the CRISPaint toolbox by developing a series of universal donor vectors to support biallelic degron tagging at the C-terminus of any gene. We show the utility of this system in human cancer cells, mouse embryonic stem cells and mouse colonic tumour organoids.

## Results

### Conditional degron-tagging using a dual fluorescent reporter system facilitates biallelic knock-in

To adapt the CRISPaint method (Figure 1A) to achieve biallelic degron tagging, we designed pairs of CRISPaint donor vectors which contained the frame selector sgRNA spacer sequences, immediately upstream from the dTAG (FKBP12^F36V^) inducible degron ^8^, fused via a short flexible linker peptide to either mClover3 or mScarleti3 (Figure 1B). When transfected together with plasmids encoding target selector and matched frame selector sgRNAs and Cas9, the donor template is linearised at a sequence encoding a glycine-rich linker immediately upstream, and the linear molecule is then integrated at the genomic locus cleaved by the target selector to generate in-frame tag fusions coupled to a generic 3’UTR. Because larger tags can cause greater perturbations to endogenous protein function in some cases, we also made versions in which the fluorescent protein markers were fused via a peptide-skipping P2A sequence to liberate the fluorescent protein during translation, leaving a relatively small tag (∼*15*KDa vs ∼*42*KDa); (Figure 1B). Of note: Schmid-Burgk et al previously published a list of target-selector sgRNA sequences suitable for engineering tag fusion at the C-terminus of any human or mouse protein ^19^.

**Figure 1:**
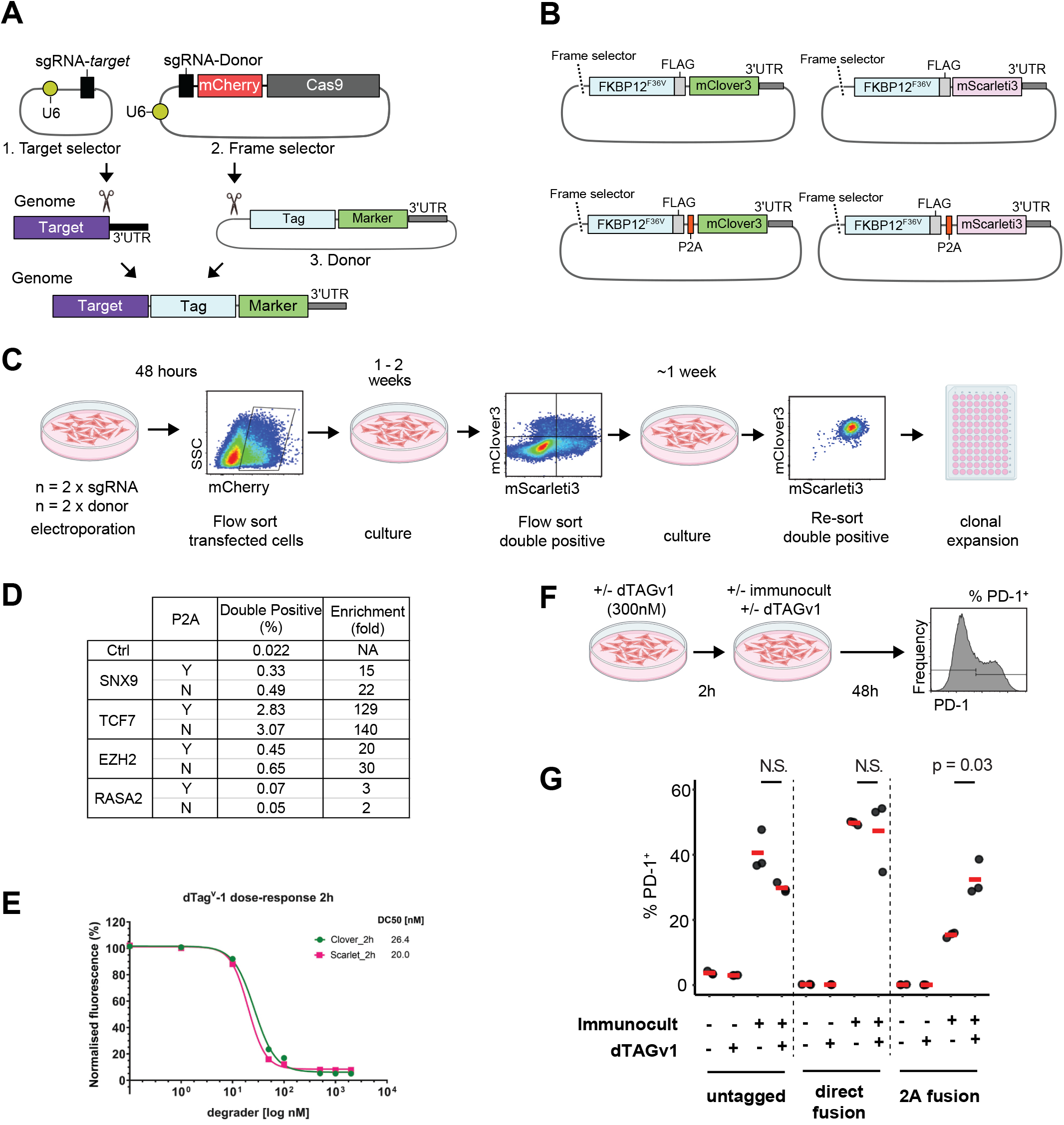
Derivation of cell lines with biallelic degron tag insertions via CRISPaint. (**A**) Schematic illustrating the principle underlying the CRISPaint approach for tagging via NHEJ. The target selector guide cleaves the target gene immediately upstream from the stop codon. Based on the frame in which the target gene is cleaved (0, 1 or 2), one of three ‘frame selector’ sgRNA plasmids is used to linearise the donor plasmid in the same frame. This increases the likelihood that the donor will be integrated in frame to generate a functional fusion protein. Note that the entire donor plasmid is typically integrated, including the 3’UTR. (**B**) Schematic shows a series of FKBP12^F36V^ donor plasmids designed specifically for biallelic degron tagging in this study. (**C**) Schematic showing the process used to derive Jurkat cells with biallelic dTAG fusions. (**D**) Table showing the fraction of cells in the mClover3 / mScarleti3 double positive gate one week after electroporation with reagents to target four different genes, using either direct or P2A-based donor plasmids. (**E**) Dose response curves show degradation of RASA2 fusion proteins across different concentrations of dTAGv1 PROTAC. (**F**) Schematic shows the experimental workflow to measure the surface expression of T cell activation marker PD-1 following activation of Jurkat cells in the presence versus absence of RASA2 degradation. (**G**) Dot plots show the percentage of cells in the PD-1^+^ gate in the presence versus absence of TCR stimulation (immunocult) and RASA2 degradation (dTAGv1). P value indicates 2-tailed t-test. N.S. = not significant at p < 0.05.

To test this approach, we first targeted four genes in Jurkat cells, which have a near-diploid genome, using pairs of donor plasmids either with or without the P2A sequence. Donor plasmids were electroporated together with target-selector (Table S1) and matched frame selector sgRNAs to integrate tags upstream from the stop codon of four genes, *RASA2, SNX9, EZH2* and *TCF7*. The frame selector sgRNA plasmid also contained mCherry, enabling FACS-purification of transfected cells. Successfully transfected cells were FACS purified 48 hours after electroporation, then returned to culture for 7 days before re-analysis (Figure 1C). Notably, in all eight cultures, we observed a larger fraction of cells in the mClover3/mScarleti3 double positive gate compared to the negative control lacking target-specific sgRNA (Figure 1D). The level of enrichment ranged from 2- to 140-fold across conditions, with relatively similar levels observed between direct and P2A-mediated fusions at the same locus (Figure 1D). We flow sorted double positive cells from RASA2 fusions and expanded the pool before re-analysis and single cell cloning. Biallelic in-frame tag fusion was confirmed by allele-specific PCR and Sanger sequencing.

We next tested the efficiency of RASA2 degradation via treatment of biallelic tagged clones with the dTAGv1 PROTAC, measuring loss of fluorescence over a 2-hour dose response. Degradation was highly efficient, removing >90% of protein expressed from each allele after 2 hours of dTAGv1 treatment above concentrations of 100nM (Figure 1E).

RASA2 was recently identified in a CRISPR KO screen as a negative regulator of T cell activation ^22^. To validate this finding, we tested the effect of RASA2 tagging and degradation on surface expression of a T cell activation marker (PD1) following stimulation of Jurkat cells in Immunocult media (Figure 1F). Consistent with published data, degradation of RASA2:dTAG (i.e., where the fluorescent protein markers were liberated via the P2A peptide) led to a significantly higher fraction of cells expressing PD-1 following Immunocult activation (Figure 1G). However, the same effect was not observed in cells where the fluorescent protein was fused directly. Here, a similarly high degree of activation occurred irrespective of dTAG treatment (Figure 1G), suggesting that the larger tag may have already disrupted RASA2 function in the non-induced state. Altogether, these data support the role of RASA2 as a negative regulator of T cell activation, and suggest that the size of C-terminal tag fusions on the RASA2 protein can negatively impact protein function, even in the absence of the degradation stimulus.

### Antibiotic selectable markers enable biallelic tagging in mouse embryonic stem cells

We next attempted the same strategy to degron-tag the mouse *Dcp2* gene, a core component of the mRNA decapping complex involved in the main pathway for cytoplasmic RNA decay ^23^, in mouse embryonic stem cells. However, we were unable to isolate double positive cells following transfection with dTAG;P2A:mClover and mScarleti3 constructs and corresponding target and frame selector guides, due to weak signal and spectral overlap between mScarleti3 and mCherry markers.

We therefore modified the selection system by creating new donor plasmids expressing drug-resistance genes. In the new donor plasmids, the fluorescent markers were replaced by Hygromycin resistance (HygroR) and Puromycin resistance (PuroR) cassettes (Figure 2A). To test this approach, mESCs were transfected with the two new donor plasmids, together with target and frame selector guide plasmids, and placed under dual antibiotic selection five days post-transfection. Surviving cells were pooled and expanded for genotyping by PCR. Modified cells were screened using primers sets designed to detect wild-type (WT), dTAG-FLAG;P2A:HygroR- and dTAG-FLAG;P2A:PuroR-tagged *Dcp2* alleles (Figure 2B). We detected correct insertion for both HygroR and PuroR tags (Figure 2C, left panel), confirming some cells within the pooled population carried at least one tagged *Dcp2* allele, while others retained a WT allele. To select cells containing both *Dcp2* alleles tagged, we obtained single-cell clones from the pooled population by limiting dilution in the presence of hygromycin and puromycin. We recovered and genotyped a total of 37 single-cell clones. Out of these, 32% (12/37) were found to have biallelic on-target tag integration, and a further 45% (17/37) showed evidence of monoallelic targeting (Figure 2D). We selected three clones (C1, C2 and C3) and confirmed by Sanger sequencing that insertions were in-frame in each case, (Figure 2C, right panel and 2E). Together, these results confirmed that the antibiotic-based CRISPaint system is an efficient method to generate and select biallelic edited mESCs.

**Figure 2.**
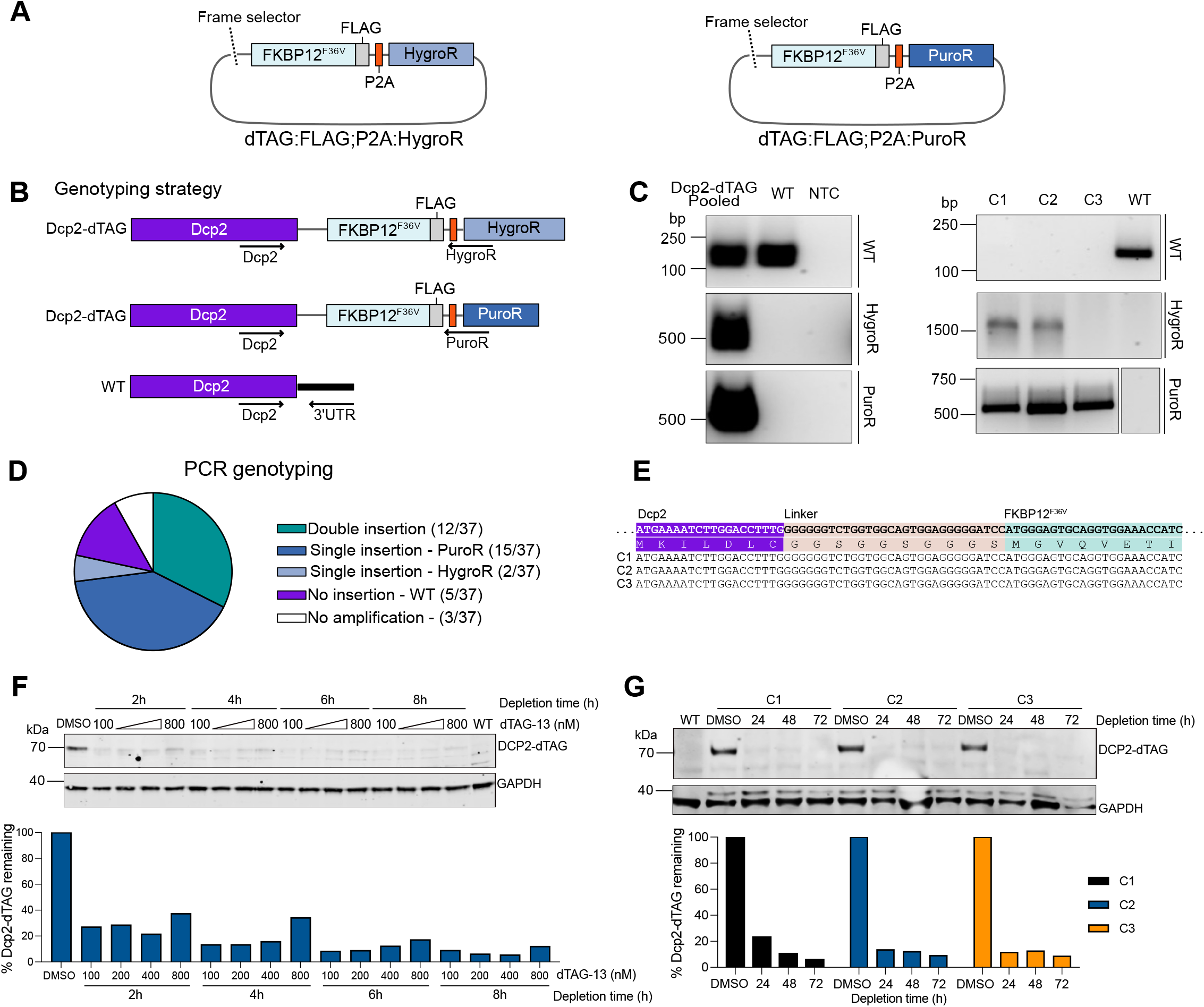
Biallelic tagging of endogenous *Dcp2* in mESCs using the CRISPaint system. **(A)**Schematic of new FKBP12^F36V^donor plasmids for the expression of hygromycin and puromycin resistance genes. (B) PCR-based genotyping strategy for screening CRISPaint-modified cells. Primer sets were designed to detect the WT (*Dcp2*) allele and the dTAG-FLAG;P2A*-*HygroR*-* (HygroR) and dTAG-FLAG;P2A-PuroR*-*tagged (PuroR) alleles. A common forward primer withing the *Dcp2* gene was used for all reactions. (**C**) PCR genotyping of *Dcp2-*dTAG pooled cells and selected single-cell clones using primer sets described in (B). HygroR and PuroR primers amplified specific bands in pooled population and single-cell clones, indicating successful insertion. (**D**) Frequency distribution of single-cell clones obtained by using plasmids in (A). (**E**) Sanger sequencing results for selected single-cell clones (C1-3), showing correct in-frame insertion. (**F**) Western blot analysis of dTAG-13-mediated depletion of DCP2-dTAG in modified mESCs. Single-cell clone C2 was treated with increasing concentrations of dTAG-13 (100, 200, 400 and 800nM) for 2, 4, 6 or 8 hours, or with DMSO as a control. WT mESCs were included as negative control for fusion protein expression. DCP2-dTAG was detected using a mouse anti-FKBP12 antibody, with GAPDH as loading control (top). Depletion efficiency was quantified by normalising DCP2-FKBP12 band intensity to GAPDH and expressing values relative to DMSO-treated cells (set to 100%) (bottom). (**G**) Western blot analysis of long-term DCP2-dTAG depletion following dTAG-13 treatment. Single-cell clones C1, C2 and C3 were treated with 250nM dTAG-13 or DMSO as a control, for 24, 48, or 72 hours. Negative control, detection antibodies and depletion levels quantification were as in (F).

Next, we assessed the expression of the DCP2-dTAG fusion protein and its depletion after dTAG-13 treatment over short periods of time. For this, we selected one single-cell clone (C2) and treated the cells with increasing concentrations of dTAG-13 (100 to 800nM) for 2, 4, 6 or 8 hours. DCP2-dTAG protein levels were analysed by western blot with an antibody against FKBP12. The levels of DCP2-dTAG protein (predicted MW size ∼ 70KDa) decreased approximately 70% within the first 2 hours of treatment with as little as 100nM dTAG-13. Protein levels kept decreasing in a time-dependent manner, with >90% depletion after 8 hours of treatment (Figure 2F). All three selected clones behaved similarly over a longer time course. Clones 1, 2 and 3 were treated with 250nM dTAG-13 for 24, 48 or 72 hours, and DCP2-dTAG protein levels were analysed by western blot. dTAG treatment resulted in an approximately 85% reduction of the fusion protein levels compared with the DMSO control, and the depletion was maintained across the 72-hour treatment (Figure 2G).

### CRISPaint degron systems allow for efficient endogenous tagging in organoids

The use of NHEJ-mediated knock-ins has dramatically increased the efficiency of endogenous protein tagging in 3D organoid models ^20^, but homozygous tagging with conditional degrons is not yet widely deployed. We utilised a mouse model of colorectal cancer with quadruple mutations in *Apc, Kras, Trp53* and *Smad4* (AKPS) ^24,25^, aiming to degrade the splicing factor SRSF1 using the second generation auxin-inducible degron system (AID2).

In addition to the degron tag fusion, AID2 requires co-expression of the auxin receptor protein from *Oryza sativa* (OsTIR1^F74G^, hereafter referred to as ‘TIR1’)^10,11^. Exposure of cells to the auxin derivative compound 5-phenyl indole-3-acetic acid (5-Ph-IAA) then enables rapid degradation of proteins tagged with a 68 amino acid peptide from *Arabidopsis thaliana* IAA17. We therefore began by generating a targeting construct for insertion of *TIR1* into the *Rosa26* safe harbour locus. This included the CAG promoter and TIR1 coding sequence separated by a lox-Neo-STOP-lox cassette to first enable selection of Neo^+^ cells using geneticin, then activation of TIR1 expression via recombination of loxP sites following exposure to Cre recombinase (Figure 3A). The targeting construct was electroporated together with a second plasmid expressing spCas9 and an sgRNA targeting the *Rosa26* locus. Single organoid clones were selected with geneticin, expanded, and genotyped, then correctly targeted clones were infected with Adenovirus expressing Cre recombinase to remove the lox-Neo-stop-lox cassette.

**Figure 3.**
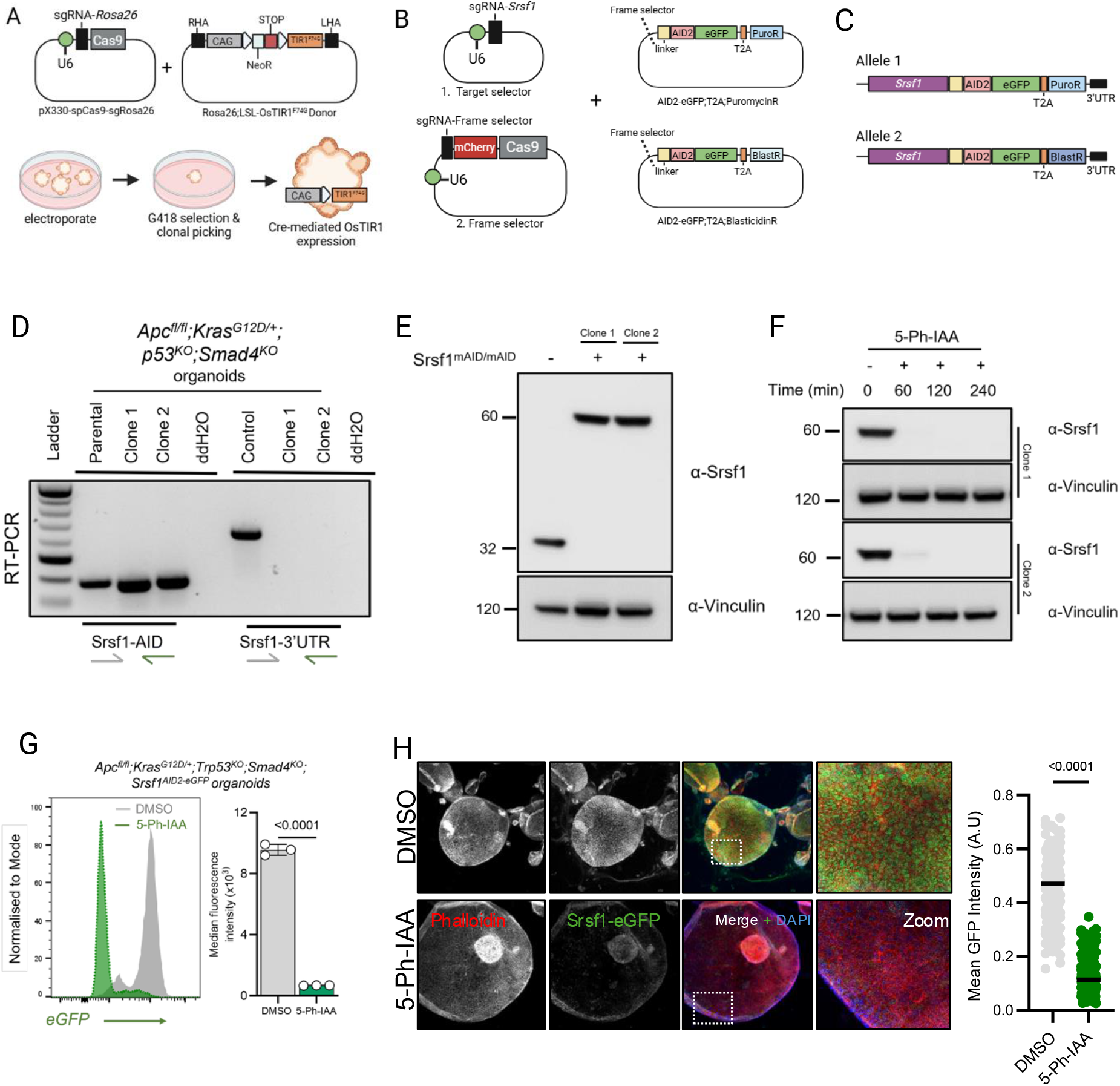
Biallelic tagging of endogenous *Srsf1* in mouse organoids. **(A)** Schematic of Rosa26 targeting strategy through HDR with Rosa26-CAG-loxP-FNF-bpA-3pA-loxP-OsTIR^F74G^ and pX330-sgRosa26-spCas9 plasmids in mouse AKPS organoids. **(B)** Schematic of endogenous *Srsf1* targeting strategy using NHEJ-AID2-eGFP plasmids with antibiotic selectable markers in AKPS;Rosa26-OsTIR^F74G^ organoids. **(C)** Schematic of endogenously tagged *Srsf1* alleles with linker-AID2-eGFP;P2A PuroR or BlastR following successful targeting. **(D)** PCR genotyping of AKPS parental line and AKPS;Srsf1^AID^ clones 1 and 2 for Srsf1-AID mutant allele and Srsf1-3’UTR wild-type allele. (E) Western blot for Srsf1 in AKPS parental line and AKPS;Srsf1^AID^ clones 1 and 2. AID-eGFP clones 1 and 2 indicate an approximate 30kDa shift consistent with successful tagging. Vinculin is used as a loading control **(F)** Western blot for Srsf1 in AKPS;Srsf1^AID^ clones 1 and 2 organoid lines following 200nM 5-Ph-IAA treatment for indicated time points of 60, 120 and 240 mins. 0 hour is untreated control. Vinculin is used as a loading control. **(G)** Flow cytometry analysis of AKPS;Srsf1^AID^ organoids following treatment with 200nM 5-Ph-IAA for 4 hours. Green histogram indicates 5-Ph-IAA treated organoids, grey histogram is DMSO treated. Median fluorescence intensity (x10^3^) is quantified. (n=3, p-value <0.0001, unpaired t-test). **(H)** Immunofluorescence images of AKPS;Srsf1^AID^ organoids treated with 200nM 5-Ph-IAA for 4 hours and visualised for Phalloidin (F-actin) and Srsf1-eGFP with nuclei stained with DAPI. Mean GFP intensity was calculated per cell using CellProfiler (n=185 DMSO vs n=279 5-Ph-IAA, p-value <0.0001, unpaired t-test).

We next generated plasmids for biallelic AID tagging with drug selection. These constructs were designed to generate direct C-terminal fusions of AID and eGFP, followed by a T2A element and either a puromycin or blasticidin resistance cassette (Figure 3B). We initially targeted the last exon of the endogenous *Srsf1* locus with the puromycin resistance construct, followed by clonal selection, resequencing of the non-integrated allele, and a second round of targeting with the blasticidin resistance construct (Figure 3B and 3C). Integration of the AID2 constructs at the *Srsf1* locus was confirmed by PCR (Figure 3D), which demonstrated successful bi-allelic insertion of AID2 conditional degron tags into the mouse AKPS colorectal cancer organoid line. To demonstrate in-frame knock-in and expression of the Srsf1-AID2-eGFP alleles we next performed western blotting, which demonstrated an approximate 30kDa band shift compared to the wild-type allele representative of biallelic AID-eGFP knock-in into the *Srsf1* locus (Figure 3E). Next, to test the degradation kinetics in this organoid model we performed a time-course with the AID2 ligand 5-Ph-IAA at 200nM. We found near complete degradation in both AKPS;Srsf1^AID2^ clones at 60 minutes which was maintained over 4 hours (Figure 3F). The addition of directly fused eGFP in the tagging construct also facilitated the visualisation of degradation kinetics by flow cytometry and immunofluorescence. Flow cytometry analysis demonstrated a significant reduction in the median fluorescence intensity of eGFP 4 hours following ligand addition (Figure 3G). Similarly, visualisation of GFP through immunofluorescence demonstrated a near complete degradation of the signal in the 5-Ph-IAA treated organoids after 4 hours of 5-Ph-IAA treatment (Figure 3H). These data demonstrate that the use of a NHEJ strategy for biallelic endogenous tagging with conditional degrons can also be successful in complex 3D organoid models.

### An expanded CRISPaint degron toolbox for endogenous tagging

Several other degron tagging systems have been developed in addition to dTAG and AID2. We have also generated CRISPaint-compatible donor vectors that enable C-terminal fusion of either BromoTag or the Small Molecule Assisted Shutoff (SMASh) tag ^9,13^. We direct the reader to recent reviews and technical articles for details of these systems and their relative merits ^7,26^. All the plasmids generated in this manuscript are detailed in Figure 4 and have been made available via the Addgene repository.

**Figure 4.**
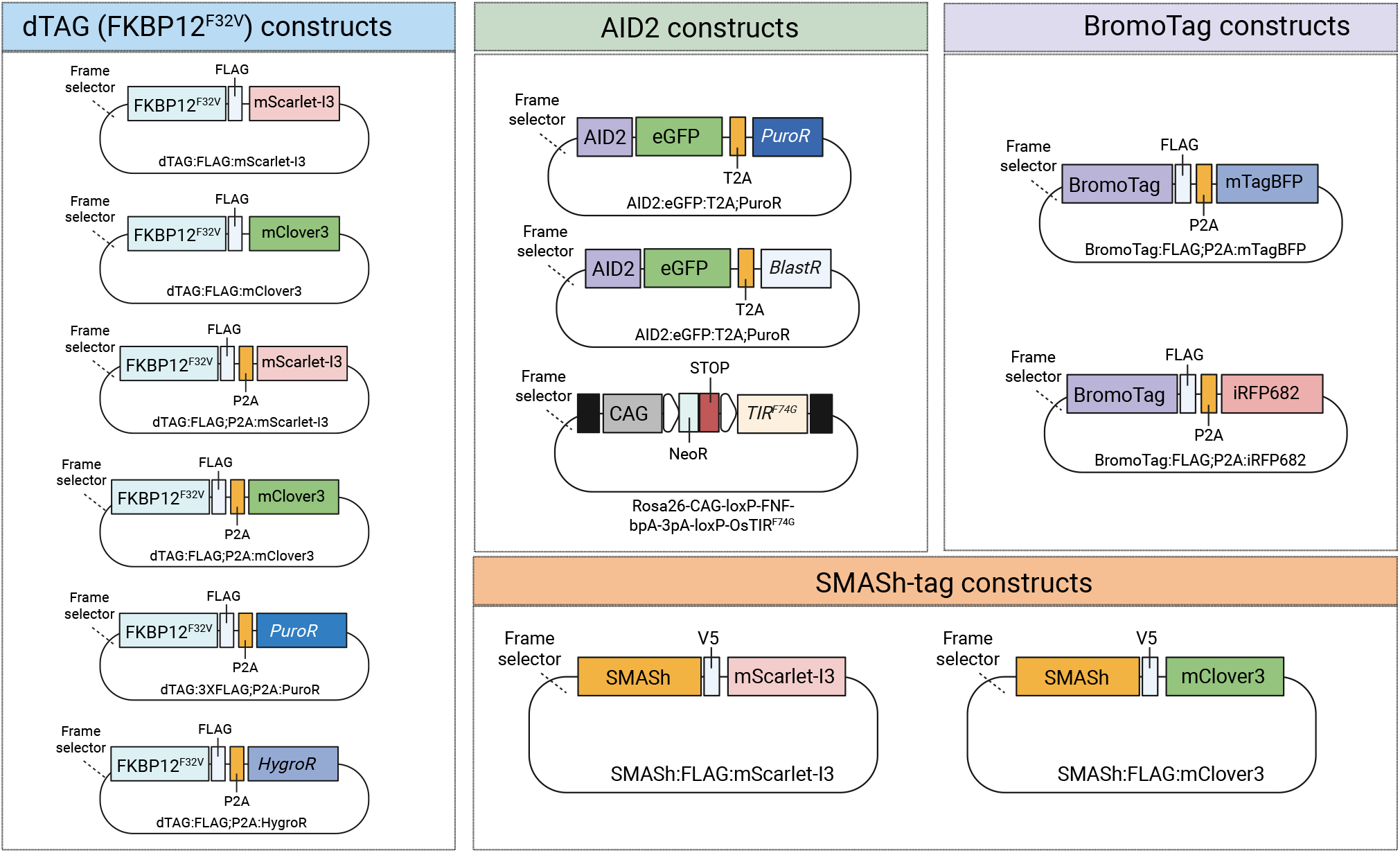
A toolbox for endogenous tagging of various degron modules. **(A)** Schematic of FKBP12^F36V^ (dTAG) donor constructs generated in this study. Each has an antibiotic or fluorescent selectable marker and FLAG epitope tag.(used in Figures 1 and 2) (**B**) Schematic of AID2 donor constructs, and Rosa26-OsTIR^F74G^ constructs generated in this study. AID2 donor plasmids contain an EGFP with a self-cleaving antibiotic resistance markers (used in Figure 3) (**C**) Schematic of BromoTag donor constructs containing mTagBFP or iRFP682 fluorescent markers. (**D**) Schematic of SMASh-tag donor constructs containing mClover3 or mScarleti3 fluorescent markers and V5 epitope tags.

## Discussion

In this paper, we have built on previously published work that combined gene editing nucleases with homology-independent integration of exogenous DNA ^17–20,27^, to develop a facile system for biallelic C-terminal fusion of inducible degron tags. By adapting the CRISPaint method, specifically, which uses a “frame selector” system to promote formation of in-frame tag fusions ^19,20^, we constructed novel targeting plasmids with dual selection markers based on either fluorescent proteins or drug selection, and show that they enable isolation of clonal populations of JURKAT lymphoma cells, mouse embryonic stem cells and colonic tumour organoids in which endogenous protein targets can be degraded >90% in under two hours following drug treatment.

Several previous publications have described methods for the genetic engineering of degron tagged cell lines, but in most cases, they have deployed homology-directed pathways of DNA repair to achieve targeted knock in ^14,15,28–30^. The activity of DNA repair pathways underlying homology-dependent DNA integration are highly variable across cell types ^16^, and require generation of bespoke HDR donor templates tailored to each integration site. In contrast, the donor templates described here are universal, meaning that only sgRNAs need to be generated before tagging a new locus, making it more straightforward to compare strategies involving different tags and selection markers for the same gene ^26^.

Despite the convenience and potential scalability of the current approach, several limitations are noteworthy. In particular, the integrated sequence includes a generic 3’UTR from the bovine growth hormone gene downstream from the stop codon, resulting in loss of any regulatory information contained in the endogenous sequence. The tagging protocol is currently optimised for C-terminal tag fusion, although adaptation for N-terminal sites of tag fusion should be feasible ^31^. Lastly, the method is also subject to challenges common to all protein tagging applications, e.g. the potential that tag fusions disrupt wildtype protein function in the non-induced state ^32–34^, and that the protocol for recombinant cell selection requires the target gene to be expressed at reasonable levels. In summary, our work combines foundational methodologies in genome engineering ^17–19^ and targeted protein degradation (^8,9,11,13^) to enable more straightforward isolation of cells carrying degron tags on both alleles. This addresses a critical bottleneck for experiments involving targeted protein degradation, enabling more tagged cell lines to be generated using fewer resources.

## Methods

### Generation of donor template plasmids

For generation of p-AID2-eGFP;T2A-PuroR and p-AID2-eGFP;T2A-BlastR a BamHI-GGGGS-AID2-eGFP;T2A-BlastR-NotI or BamHI-GGGGS-AID2-eGFP;T2A-PuroR-NotI DNA construct was synthesised by Twist Biosciences. These constructs were digested with BamHI and NotI before being ligated into digested backbone of pCRISPR-HOT_tdTomato (Addgene: 138567). Positive clones were then screened by restriction digest and Sanger sequencing. dTAG:FLAG:mClover3/dTAG:FLAG:P2A:mClover3 and dTAG:FLAG:mScarleti3/dTAG:FLAG:P2A:mScarleti3 fragments were synthesised by Twist Bioscience with a 5’ BamHI and 3’NotI site, which was used to replace the equivalent BamHI/NotI fragment as described above. The dTAG:FLAG;P2A:HygroR and dTAG:FLAG;P2A:PuroR plasmids were generated by replacing fluorescent markers with hygromycin (HygroR) and puromycin (PuroR) cassettes. HygroR and PuroR genes were PCR-amplified from previously established plasmids using specific primers to amplify the full resistance cassettes while introducing HindIII and AgeI restriction sites at the 5’ and 3’ ends, respectively. The mClover3 donor plasmid was PCR-amplified to introduce corresponding restriction sites and remove mClover3 marker. PCR products (HygroR, PuroR and donor backbone) were purified, digested with HindIII and AgeI, and ligated using T4 DNA ligase. Correct clones were selected by restriction digestion and Sanger-sequenced.

To generate Rosa26-CAG-loxP-FNF-bpA-3pA-loxP-OsTIR^F74G^ plasmid, OsTIR^F74G^ was amplified by PCR using Phusion polymerase from pAAV-hSyn-OsTIR^F74G^ (Addgene: 140730) with 5’-AscI and 3’-PacI restriction sites. Digested OsTIR^F74G^ PCR product was then ligated into AscI and PacI digested Rosa26-CAG-loxP-FNF-bpA-3pA-loxP-tdTomato backbone (Addgene: 180152). Positive clones were then screened by restriction digest and PCR amplification of the OsTIR^F74G^ insert. Each donor plasmid has been submitted to Addgene under submission ID 87793.

### Generation of tagged Jurkat cell lines

Jurkat cells were routinely cultured in RPMI with 10% FBS and 1% Pen/Strep. The following target-specific sgRNA spacer sequences were used RASA2: TAAGATGCTTTCCCAACAAT; EZH2: GAGGAGGTAGCAGATGTCAA; TCF7: AACCAGCAGACGGATTGGTG; SNX9: CAGCCGCTTTCCAGTGATGT. Electroporation of CRISPaint plasmids was conducted using a Neon NxT Electroporation system. An equal mass of each plasmid (3 μg total) and 1×10^6^ cells were resuspended in Resuspension R Buffer at a final density of 1×10^7^ cells/mL, aspirated into a Neon NxT 100uL tip, and electroporated with 3 pulses at 1325V and 10 ms pulse width, before plating into a pre-warmed 6 well plate. After 48 hours, cells were sorted for mCherry positive (transfected) cells, and at 7 days post transfection, the percentage of cells expressing both mScarleti3 and mClover3 was quantified on a Beckman Coulter Cytoflex flow cytometer. To isolate *RASA2* homozygous tagged cells, the double positive population was FACS purified, pooled, cultured for a further 2 weeks, then individual double positive cells were re-sorted into pre-conditioned media in a 96 well plate, and single cell clones were expanded and genotyped via PCR and Sanger sequencing of insertion junctions.

For TCR stimulation, 6 × 10^6^ cells were seeded. The following day, the culture was split into four tubes, spun down, then each tube was resuspended in 2955 μl media containing either 300 nM dTAGv1 or DMSO control. After two hours of dTAGv1 exposure, 45μl Immunocult human CD3/CD28 T Cell Activator solution (Stem Cell Technologies), or an equivalent volume of media, was added. After 48 hours of incubation, cultures were centrifuged at 400g for 5 minutes, incubated with DAPI

(1mg / ml in FACS buffer) for 15 minutes, spun and resuspended in 25 μl FC block buffer (BD Pharmingen) and incubated at room temperature. After 10 minutes, 25 μl anti-PD1 Brilliant Violet 421 antibody (BD) was added to a final working dilution of 1:100, and cells were incubated at 4C in the dark. After 30 minutes, 200 μl of FACS buffer was added, cells were spun, supernatant was removed, and cells were resuspended in 100 ul Cytofix buffer (BD) and incubated at 4C. After 10 minutes, cells were spun down, resuspended in 100 ul PBS and analysed on a Beckman Coulter Cytoflex flow cytometer.

### Emrbyonic stem cell culture and degron induction

v6.5 mouse embryonic stem cells (mESCs) were cultured in StableCell high glucose Dulbecco’s modified Eagle Medium (DMEM, Sigma), supplemented with 15% heath-inactivated foetal calf serum (FCS) (Gibco), 100 U/ml penicillin-streptomycin, 100U/ml leukemia inhibitory factor (LIF; Stemcell technologies), 1X sodium pyruvate (Gibco), 1X non-essential amino acids (Gibco) and 40 µM 2-mercaptoethanol (Gibco). Cells were maintained on 0.1% gelatine-coating plates and incubated at 37°C in a humidified 5% CO_2_ atmosphere.

A guide RNA (gRNA) targeting *Dcp2* (5’*-*CAATACTCTTGCTGGCTCAA-3’) was cloned into the target selector plasmid using BbsI restriction sites. mESCs were grown overnight and transfected at ∼60-70% confluency with antibiotic-resistance markers (1µg DNA of each plasmid) and Lipofectamine 2000 according to manufacturer’s instructions in 6-well plates. Cells were placed under dual antibiotic selection (250µg/ml Hygromycin (Invitrogen), 0.5µg/ml Puromycin (Merck)) at 5 days post-transfection. The medium was replaced daily for five days, and the cells were left to recover. Antibiotic resistant cells were pooled and genotyped by PCR and further diluted (1cell/well) to isolate single-cell clones. Platinum Direct PCR Universal Master Mix (ThermoFisher) was used for all screening PCRs of CRISPaint-modified mESCs, following manufacturer’s instructions. Genomic DNA was prepared by resuspending 5 × 10^5^ cells in 40 µl of lysis buffer supplemented with proteinase K, followed by heat inactivation at 98°C for 1 minute and centrifugation at maximum speed to pellet debris. 2 µl of the supernatant were used for each PCR reaction. Three primer sets were used to screen for WT and dTAG-tagged *Dcp2* alleles (**Table 1**).

**Table 1.**
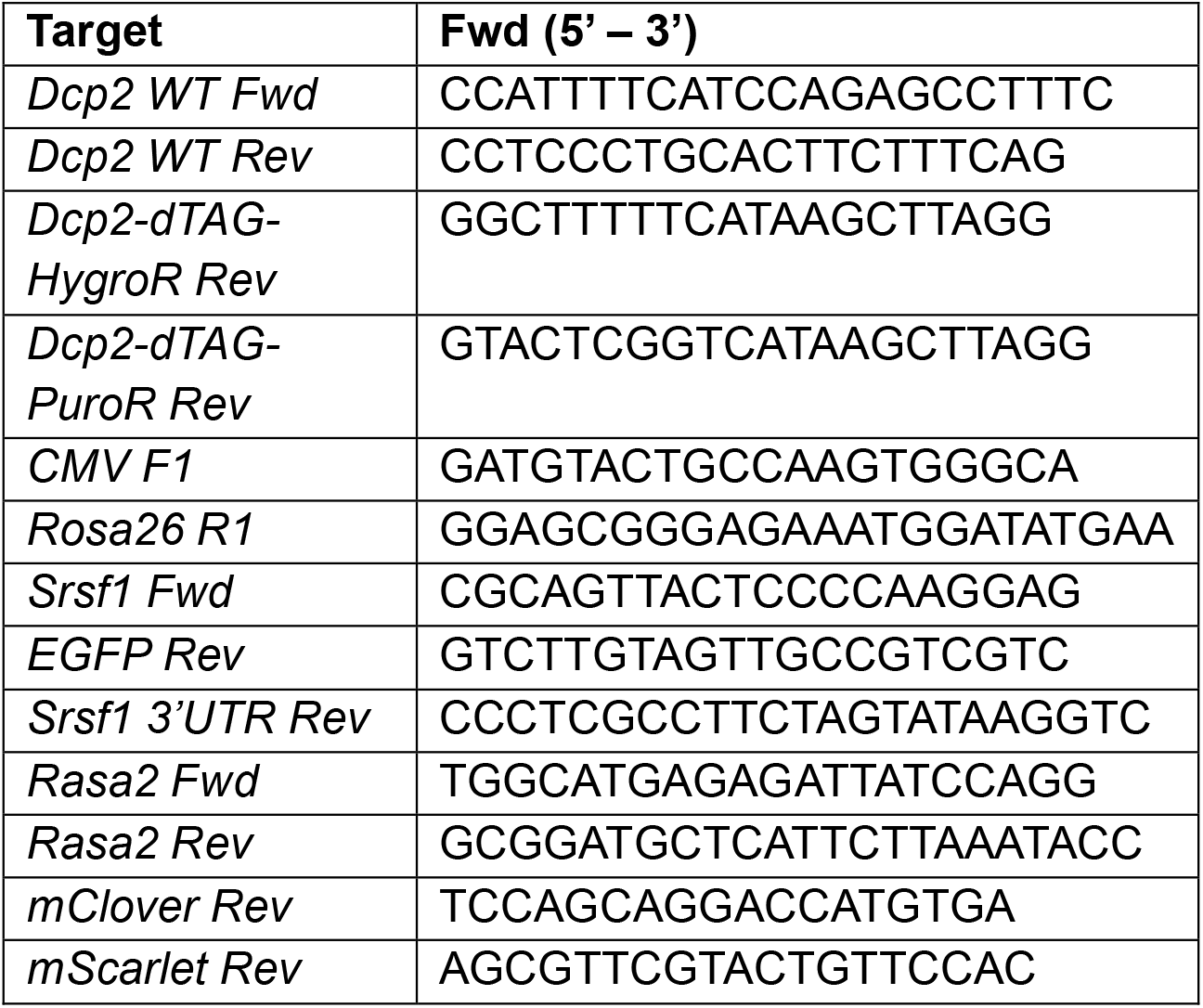
List of primers used for PCR screening.

For protein degradation, tagged mESCs were plated in 24-well plates and allowed to grow overnight. Cells were washed once with 1x PBS and incubated in fresh medium containing DMSO or increasing concentrations of dTAG-13 (Tocris) (100nM, 200nM, 400nM and 800nM). Cells were treated for 2, 4, 6 or 8 hours, after which cell lysates were collected for western blot analysis. For long-term depletions experiments, WT mESCs and single-cell clones C1, C2 and C3 were plated in 6-well plates for overnight growth. Cells were then incubated in fresh medium with DMSO or 250nM dTAG-13 for 24, 48 or 72 hours.

### Generation of Apc^fl/fl^;Kras^G12D/+^;Trp53^KO^;Smad4^KO^ mouse organoid tagged lines

Apc^fl/fl^;Kras^G12D/+^;Trp53^KO^;Smad4^KO^ organoids were embedded in phenol-red free Basement Membrane Extract (BME, Cultrex, R&D Systems) and cultured in Advanced DMEM/F12 (Gibco) supplemented with L-Glutamine, HEPES and Primocin (InvivoGen), supplemented with 1X B27 (ThermoFisher Scientific), 1X N2 (ThermoFisher Scientific), 50ng/mL EGF (Peprotech) and 1% Noggin conditioned media (described in^29^).

Apc^fl/fl^;Kras^G12D/+^ organoids were electroporated with sgRNAs targeting *Trp53* and *Smad4* following the protocol described in ^35^. sgRNAs were cloned into pX330-U6-Chimeric_BB-CBh-hSpCas9 plasmid (Addgene #42230). Briefly, 1×10^6^ single cells were electroporated using NEPA21 (Nepa Gene) in Opti-MEM media before being cultured in antibiotic-free media. After 7 days recovery organoids were digested to single cells and selected with 20 µM Nutlin-3 (Selleck Chemicals) and 20 ng/mL TGFβ1 (R&D Systems). Following this, single organoid clones were collected and expanded for confirmation of Trp53/Smad4 knockout by western blotting. sgRNA guide sequences are as follows: *Trp53:* 5’-CACCGCCTCGAGCTCCCTCTGAGCC-3’, *Smad4:* 5’-CACCGCCAGGACAGCAGCAGAA-3’.

### Generation of Rosa26-OsTIR1 and Srsf1^AID^ mouse organoid lines

Apc^fl/fl^;Kras^G12D/+^;Trp53^KO^;Smad4^KO^ were generated as previously described ^24,25^. To generate AKPS;Rosa26-LSL-OsTIR1^F74G^ organoids, 1×10^6^ AKPS cells were electroporated with 5 µg pU6-sgRosa26-1_CBh-Cas9-T2A-BFP and 7.5 µg of Rosa26-CAG-loxP-FNF-bpA-3pA-loxP-OsTIR^F74G^ following a previously described protocol ^35^. Briefly, single cells were electroporated using NEPA21 electroporator (Nepa Gene) in Opti-MEM media before being cultured in antibiotic-free advanced DMEM/F12 (Gibco), containing 1X B27 and N2, with 50ng/mL EGF (Peprotech), 1% Noggin conditioned-media and supplemented in 10µM Y27632. 4 days following electroporation cells with positive integration were selected with 500ug/mL G418/Geneticin (ThermoFisher Scientific). Geneticin resistant clones were then manually picked and expanded. Correct integration was confirmed via PCR of genomic DNA using primers CMV F1 and Rosa26 R1 (Table 1), followed by Sanger sequencing.

For Srsf1 targeting, AKPS;Rosa26-LSL-OsTIR1^F74G^ organoids were electroporated as described above with the following plasmids to allow endogenous targeting of exon 4 of Srsf1 (ENSMUSG00000018379): 5 µg pSPgRNA-sgSrsf1 (Fwd: 5’-GAGCGAGATCTGCTATGACG-3’), 5µg pCas9-mCherry-Frame+0 (Addgene: 66939) and 7.5µg p-AID2-eGFP;T2A-BlastR. 4 days following electroporation cells with positive integration were selected with 10ug/mL Blasticidin (Gibco). Correct integration was confirmed via PCR of genomic DNA using primers Srsf1 Fwd and EGFP Rev (Table 1), followed by Sanger sequencing. The Srsf1 wild-type allele was amplified using Srsf1 Fwd and Srsf1 3’UTR Rev (Table 1). A 2bp deletion in the wild-type Srsf1 allele allowed retargeting with 5 µg pSPgRNA-sgSrsf1 (Fwd: 5’-TGCTGACGGGGAGAATAGCG -3’), 5µg pCas9-mCherry-Frame+2 (Addgene: 66941) and 7.5µg p-AID2-eGFP;T2A-PuroR. 4 days following electroporation cells with positive integration were selected with 2ug/mL Puromycin (Gibco), and clonal organoids were confirmed for correct integration via western blot and Sanger sequencing.

5-Ph-IAA (Tocris) in DMSO was used at 200nM for indicated time points throughout the experiments.

### Western blotting

For western blot analysis, wild-type and CRISPaint-modified cells were lysed in RIPA buffer supplemented with EDTA-free protease inhibitor cocktail (Roche), incubated on ice for 20 minutes and centrifuged at maximum speed for 15 minutes at 4°C. The supernatant was collected as total cell lysate. Protein concentration was measured by a BCA assay. Equal amounts of protein were resolved on 4-12% Bis-tris SDS-PAGE gels (Sigma) and transferred onto nitrocellulose or PVDF membranes. For Srsf1 and β-actin, membranes were blocked in 5% skim milk powder in 0.1% PBS-Tween20 (PBST) for 1 hour at room temperate before incubated overnight in 1:1000 Srsf1 (Abcam, ab129108) and 1:2000 Vinculin (Abcam, ab129002) in 5% skim milk powder-PBST. Anti-mouse and anti-rabbit-HRP conjugated antibodies (Cell Signalling Technology, 7076 and 7074) were incubated at 1:5000 in 3% skim milk powder-PBST for 1 hour at room temperature. Membranes were then imaged on ImageQuant8000 (Amersham−) following chemiluminescence detection. For FKBP12 and GAPDH, membranes were blocked in 5% FCS in 0.1% PBST for one hour at room temperature and probed with antibodies against mouse FKBP12 (Abcam 24373, 1:1000) and mouse GAPDH (6CS Thermo Fischer, 1:2000), followed by species-specific IRDye-conjugated secondary antibodies (LI-COR, 1:10.000) incubation. Band intensities were quantified using a LI-COR Odyssey Fc imaging system.

### Organoid Flow Cytometry

Organoids were prepared for FACS by the following protocol: organoids were digested to single cells for 20-25 minutes at 37°C in TrypLE express (+ Y27632). The reaction was stopped by adding equal volume of 1:1 volume of 0.1% BSA-PBS. Cells were the resuspended in Advanced DMEM/F12 media filtered through a pre-wet 0.45µm filter, and centrifuged for 500xg for 5 minutes at 4°C. Cells were fixed in 2% paraformaldehyde for 10 minutes on ice, washed twice with PBS and analysed on the BD Biosciences Fortessa−.

### Organoid Immunofluorescence

BME embedded organoids were washed once in PBS, fixed in 2% PFA on ice for 20 minutes, before three more washes. Cells were then permeabilised in 0.5% Triton X-100 for 20 minutes at room temperature, before washed once in IF buffer (0.2% Triton X-100, 0.05% Tween-20 in PBS). Organoids were then stained with 1:400 Phalloidin-CF568 (VWR) for 20 minutes at room temperature, before two washes with IF buffer. Mounting media with DAPI (VECTASHIELD) was then added before imaging Nikon Eclipse confocal microscope.

## Acknowledgements

We are grateful to the IGC Flow Cytometry Core and Advanced Imaging Resource for technical support. AW acknowledges funding from the Medical Research Council, UK, Award no. MC_PC_21040. SM acknowledges funding from Wellcome Trust grant 221737/Z/20/Z and a Wellcome Trust PhD fellowship to CGC 319808/Z/24/Z. KBM acknowledges funding from Cancer Research UK (CRUK) under a Career Development Fellowship (A19166 to K.B.M.) and a Small Molecule Drug Discovery Project Award (A25808 to K.B.M.), a European Research Council under Starting Grant (COLGENES–715782 to K.B.M.) and the MRC (project grant MR/X008762/1 to K.B.M.)

